# A phosphorylation-dependent partner-switching-like module controls heterocyst polysaccharide layer formation in *Anabaena* sp. strain PCC 7120

**DOI:** 10.64898/2026.04.08.717354

**Authors:** Mai Harada, Satoshi Matsuoka, Shigeki Ehira

**Affiliations:** Department of Biological Sciences, Graduate school of Science, Tokyo Metropolitan University, Hachioji, Tokyo, Japan; Department of Biochemistry and Molecular Biology, Graduate School of Science and Engineering, Saitama University, Saitama, Saitama, Japan

## Abstract

Heterocysts are specialized cells formed by filamentous cyanobacteria to enable nitrogen fixation under aerobic conditions. These cells are surrounded by a multilayered envelope, including a heterocyst-specific polysaccharide (Hep) layer that restricts oxygen diffusion and protects nitrogenase activity. Although several genes required for Hep biosynthesis have been identified, the regulatory mechanisms controlling Hep layer formation remain poorly understood. Here, we show that Hep layer formation in *Anabaena* sp. PCC 7120 is regulated by a phosphorylation-dependent partner-switching-like system. The phosphatase HenR and the STAS domain-containing protein All4160 are both required for Hep formation and nitrogen fixation. A nonphosphorylatable All4160 variant restores Hep formation and nitrogen fixation in a *henR* disruptant, indicating that phosphorylation negatively regulates All4160 function. Using a bacterial two-hybrid assay, we identified two candidate kinases, Alr3423 and All2284, that interact with All4160, and both proteins phosphorylate All4160 in vitro. Genetic analysis revealed that deletion of alr3423, but not all2284, suppresses the nitrogen fixation defect of the *henR* disruptant, suggesting that Alr3423 is a major kinase regulating All4160 in vivo. These findings uncover a phosphorylation switch that controls heterocyst polysaccharide layer formation and expand the role of partner-switching systems beyond σ factor-dependent transcription to the direct regulation of a biosynthetic enzyme involved in exopolysaccharide (EPS) synthesis.

**Importance:** Exopolysaccharides (EPS) are involved in diverse aspects of bacterial physiology, including environmental acclimation, motility, host association, and cellular differentiation, yet their regulatory mechanisms remain poorly understood. Cyanobacteria play central roles in global carbon and nitrogen cycles through photosynthesis and nitrogen fixation, the latter requiring a micro-oxic environment provided by specialized cells called heterocysts. This study reveals a previously unrecognized mechanism for controlling polysaccharide biosynthesis, in which a partner-switching-like system directly regulates the activity of a biosynthetic enzyme. By expanding the role of partner-switching systems beyond σ factor–dependent transcription, these findings provide a framework for understanding how bacteria control EPS production in response to environmental and developmental cues.

## Introduction

Cyanobacteria are a phylogenetically and physiologically diverse group of bacteria capable of oxygenic photosynthesis and play central roles in global carbon and nitrogen cycles. In many ecosystems, they function as major primary producers by fixing carbon dioxide and, in some species, dinitrogen through nitrogen fixation. Because nitrogenase, the enzyme responsible for nitrogen fixation, is highly sensitive to oxygen, cyanobacteria have evolved distinct strategies to protect this activity from oxygen inactivation (1). Unicellular cyanobacteria typically temporally separate photosynthesis and nitrogen fixation, restricting nitrogen fixation to the night, whereas many filamentous cyanobacteria spatially separate these processes. The filamentous cyanobacterium *Anabaena* sp. strain PCC 7120 (hereafter *Anabaena*) differentiates cells called heterocysts, which are specialized to nitrogen fixation, at the semi-regular interval along its filament under nitrogen-deprived conditions (2, 3). Heterocysts provide a microoxic intracellular environments that supports nitrogenase activity through multiple coordinated mechanisms. During differentiation, oxygen-evolving photosystem II is inactivated, while respiration is activated to consume residual oxygen. In addition, heterocysts are surrounded by a multilayered envelope consisting of heterocyst-specific glycolipid and polysaccharide layers (Hgl and Hep layers), which function as a barrier to oxygen diffusion. Proper formation of the Hep layer is essential for maintaining the microoxic environment in heterocysts.

Synthesis of the Hep layer depends on a cluster of genes known as the HEP island, which comprises 17 genes (alr2825 to alr2841) that are induced under nitrogen-deprived conditions (4, 5). Approximately half of genes in the HEP island encode glycosyltransferase and are required for Hep layer formation. In addition, several genes encoding glycosyltransferases located outside the HEP island, such as *hepB*, alr3699, and all4160, are also involved in the Hep layer formation (6). Genes with regulatory functions have also been implicated in Hep layer synthesis. *hepK*, which encodes a histidine kinase of a two-component regulatory system, is required for the expression of *hepA*, one of the HEP island genes (7). HepK phosphorylates the response regulator DevR; however, the function of DevR remains unclear because it lacks an apparent output domain (8). *hepN* also encodes a histidine kinase and regulates the expression of *hepA* and *hepB* (9). In addition, *hepS*, encoding a Ser/Thr kinase, and *henR*, encoding a Ser phosphatase, are required for proper formation of the Hep layer (10). However, the regulatory mechanisms underlying Hep layer biosynthesis remain poorly understood.

One potential regulatory mechanism underlying Hep layer formation involves reversible protein phosphorylation. The partner switching system (PSS) is a regulatory mechanism that controls gene expression in response to environmental changes in bacteria. This system has been most extensively studied in *Bacillus subtilis*, where it regulates the activity of σ factors by modulating their interaction with RNA polymerase, thereby controlling the expression of genes involved in sporulation and stress responses (11, 12). The PSS consists of three types of proteins: a serine kinase that functions as an anti-σ factor, a serine phosphatase containing a SpoIIE-like domain, and a protein containing a sulfate transporter and anti-sigma factor antagonist (STAS) domain, whose phosphorylation state is controlled by the kinase and phosphatase (13). In this system, phosphorylation of the STAS protein promotes the interaction between the kinase and the σ factor, resulting in transcriptional repression, whereas dephosphorylation promotes association between the STAS protein and the kinase, leading to release of the σ factor and activation of transcription. Thus, the phosphorylation state of the STAS protein determines the interaction partner of the kinase, enabling dynamic switching of protein-protein interactions that regulates σ factor activity.

Although PSS-like regulatory systems have been identified in various bacteria, their roles in cyanobacteria remain poorly understood. Notably, All4160 contains an N-terminal STAS domain, suggesting that it may participate in a phosphorylation-dependent regulatory mechanisms analogous to the PSS. In addition, HenR is predicted to function as a Ser phosphatase containing a SpoIIE-like domain and is required for proper Hep layer formation, raising the possibility that HenR and All4160 constitute a PSS-like regulatory module controlling Hep layer biosynthesis. In this study, we investigated the functional relationship between HenR and All4160, focusing on the role of phosphorylation of the All4160 STAS domain in the regulation of Hep layer formation.

## Results

### A nonphosphorylatable All4160 variant suppresses the defects of the *henR* mutant

To investigate the functional relationship between HenR and All4160, we examined whether the phosphorylation state of All4160 affects Hep layer formation and nitrogen fixation. As previously reported (10), the *henR* disruptant (Δ*henR*) generated in this study showed impaired growth under nitrogen-fixing conditions, whereas growth was comparable to the wild-type strain (WT) in the presence of nitrate (Figs. 1A and B). Consistently, Alcian blue staining revealed that Δ*henR* failed to form a proper Hep layer (Fig. 1C). Similar to Δ*henR*, the All4160 disruptant (Δ4160) also exhibited impaired diazotrophic growth and defective Hep layer formation (Fig. S1) (6). To assess the role of phosphorylation of the STAS domain of All4160, we generated a Δ*henR* strain carrying a plasmid expressing an All4160 variant in which the putative phosphorylatable serine residue was replaced with alanine (Δ*henR*/S74A) (Fig. S2). The Δ*henR*/S74A strain restored growth under nitrogen fixing conditions and formed heterocysts with a Hep layer detectable by Alcian blue staining (Figs. 1B and 1C). In contrast, expression of wild-type All4160 in the Δ*henR* background did not restore diazotrophic growth (Fig. S3).

**Figure 1.**
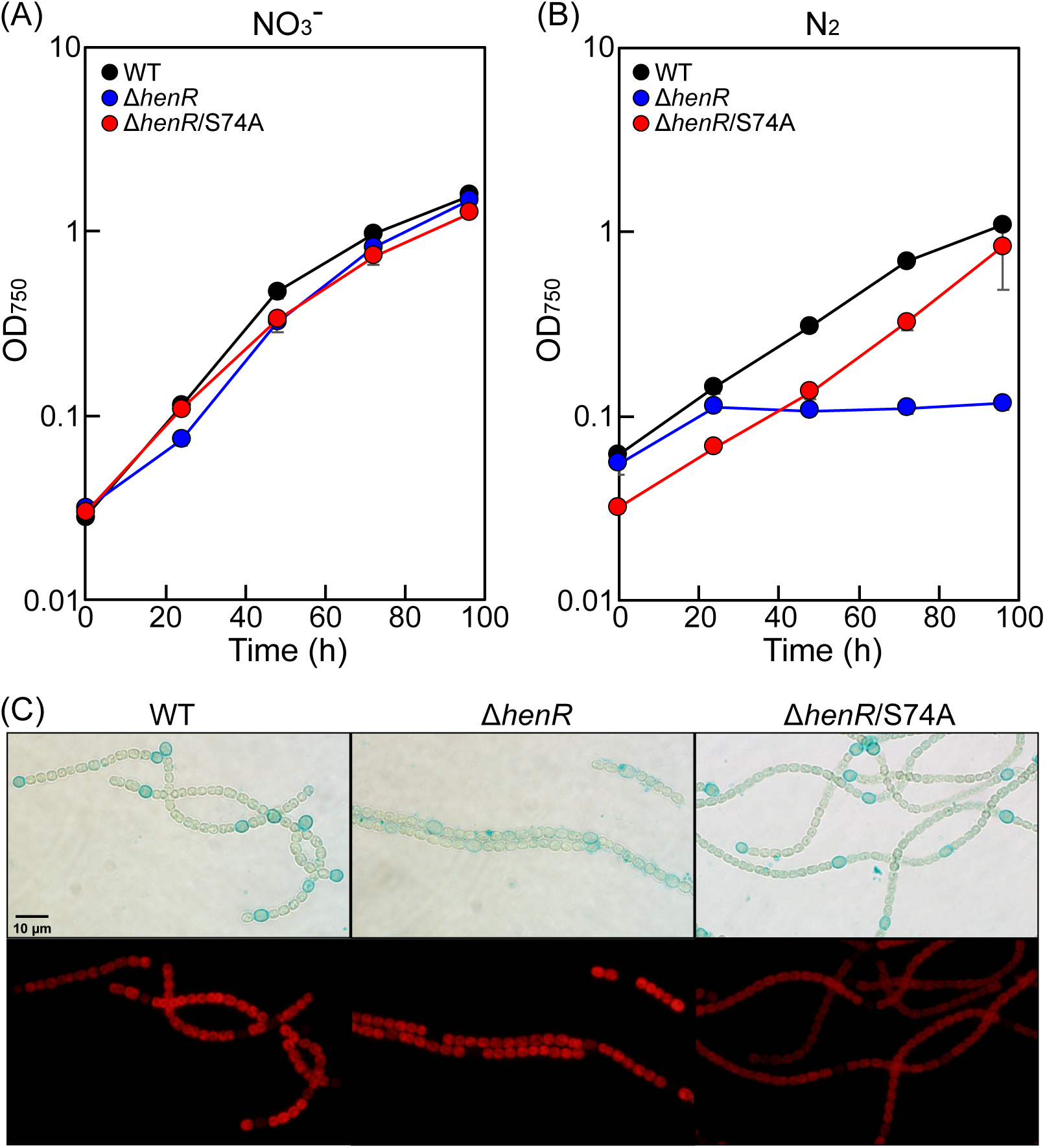
A nonphosphorylatable All4160 variant restores Hep layer formation and nitrogen fixation in the absence of HenR. Preventing phosphorylation of All4160 rescues the Hep layer defect and nitrogen fixation phenotypes of the *henR* mutant (Δ*henR*). Growth of the wild-type strain (WT) (black circles), Δ*henR* (blue circles), and Δ*henR* carrying a plasmid expressing an All4160 variant in which the phosphorylatable serine residue was replaced with alanine (Δ*henR*/S74A) (red circles) with nitrate (A) or dinitrogen (B) as nitrogen sources. (C) Micrographs of Alcian blue-stained cells of WT (left panels), Δ*henR* (center panels), and Δ*henR*/S74A (right panels). The upper panels show bright-field images, and the lower panels show autofluorescence.

To determine whether HenR affects transcription of Hep biosynthesis genes, we measured the transcript levels of *hepA* and *hepB* during heterocysts differentiation (Fig. S4). In Δ*henR*, the transcript levels of both genes were induced after 8 h of nitrogen deprivation, similarly to WT, remained high after 24 h, whereas those in WT decreased at 24 h. These results indicate that HenR does not regulate Hep layer formation at the transcript level.

Taken together, these results indicate that phosphorylation of the STAS domain negatively regulates All4160 activity in Hep layer formation.

### Identification of kinases that phosphorylate the All4160 STAS domain

We next sought to identify kinases responsible for phosphorylation of the All4160 STAS domain. The genome of *Anabaena* encodes five candidate serine kinases (All1702, All2284, Alr3423, Alr3655, and Alr3760). To test interactions between these kinases and the All4160 STAS domain, we performed a bacterial adenylate cyclase two-hybrid (BACTH) assay. Because phosphorylation of the conserved serine residue in the STAS domain weakens its interaction with kinases (14), we used a nonphosphorylatable variant in which the putative phosphorylation site (Ser74) was replaced with alanine [4160STAS(S74A)] (Fig. 2A). Qualitative analysis on MacConkey agar showed that All2284 and Alr3423 interacted with 4160STAS(S74A) (Fig. 2B). Interactions of both kinases with the wild-type STAS domain (4160STAS) were weaker than those with 4160STAS(S74A), suggesting that these kinases phosphorylate Ser74 of the All4160 STAS domain (Fig. 2C).

**Figure 2.**
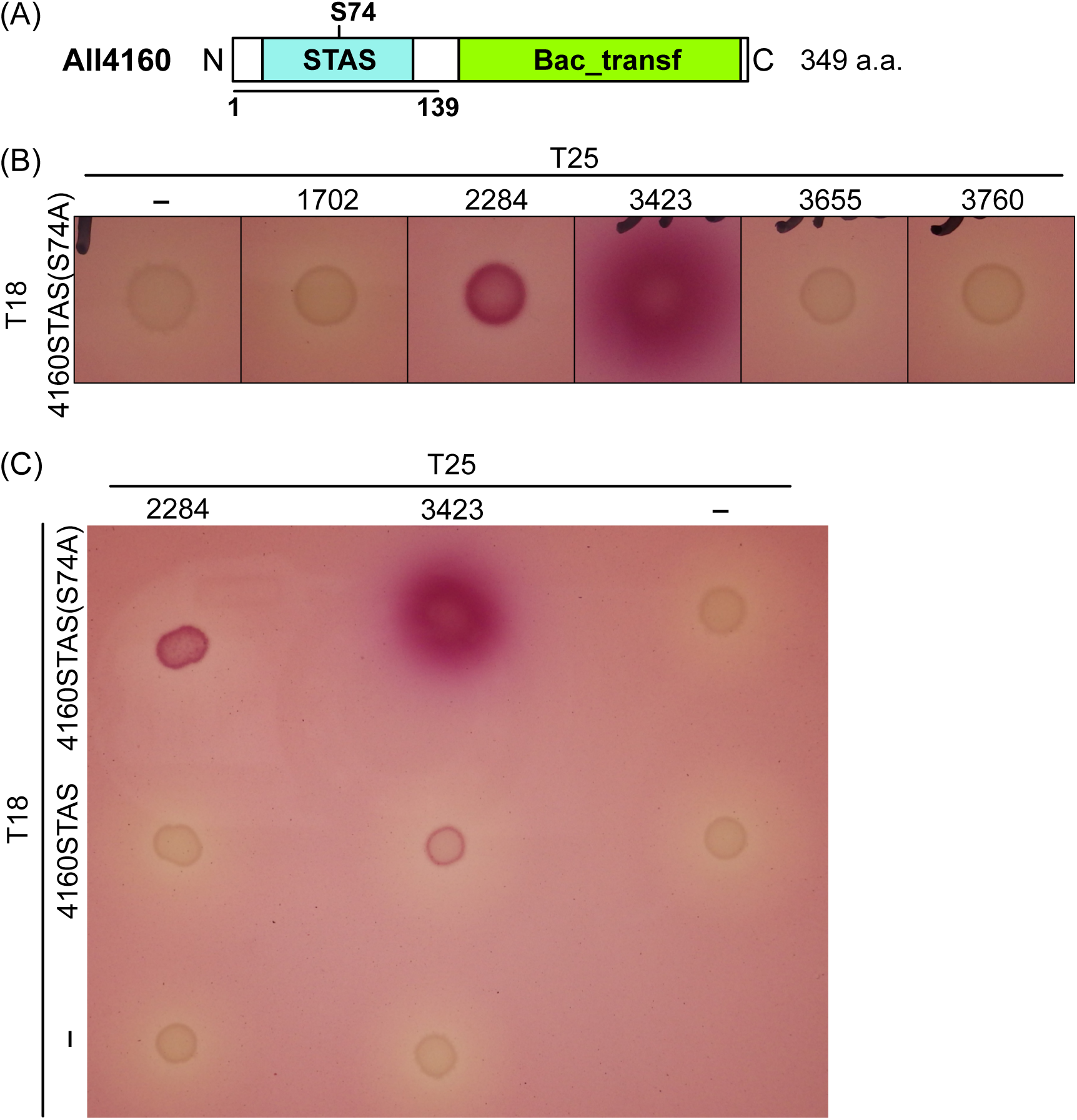
Phosphorylation–dependent interaction between the All4160 STAS domain and candidate kinases. Protein-protein interactions were analyzed using a bacterial two-hybrid (BACTH) system. All4160 interacted with two candidate kinases, All2284 and Alr3423. Interaction with these kinases was stronger when the phosphorylatable serine residue of the All4160 STAS domain was replaced with alanine (S74A). (A) Domain structure of All4160. The protein contains an N-terminal STAS domain and a C-terminal Bacterial sugar transferase (Bac_transf) domain. S74 indicates the putative phosphorylation site. The STAS region (residues 1-139) was used in the BACTH assay. (B) Screen for kinases interacting with the All4160 STAS domain (S74A variant). The STAS domain of All4160 carrying the S74A substitution [4160STAS(S74A)] was fused to the T18 fragment of adenylate cyclase, and candidate kinases (All1702, All2284, Alr3423, Alr3655, or Alr3760) were fused to the T25 fragment. The plasmids were cotransformed into *E. coli*, and interactions were detected on MacConkey agar containing maltose. “–” indicates plasmids expressing only the T18 or T25 fragments. (C) Interaction of the wild-type STAS domain with All2284 and Alr3423.

To determine whether All2284 and Alr3423 phosphorylate the All4160 STAS domain, we performed in vitro phosphorylation assays. Recombinant proteins (All2284, Alr3423, 4160STAS, and 4160STAS(S74A)) were expressed as His-tagged proteins and purified by affinity chromatography (Fig. 3A). All2284 or Alr3423 was incubated with 4160STAS in the presence or absence of ATP, and the proteins were analyzed by Phos-tag SDS-PAGE, in which phosphorylated proteins migrated more slowly than nonphosphorylated forms. When All2284 was incubated with 4160STAS in the presence of ATP, a mobility shift of 4160STAS was observed, whereas no shift was detected in the absence of ATP (Fig. 3B, lanes 5 and 6). In contrast, incubation of All2284 with 4160STAS(S74A) did not cause a band shift even in the presence of ATP (Fig. 3B, lanes 7 and 8). These results indicate that All2284 phosphorylates Ser74 of the All4160 STAS domain in an ATP-dependent manner. Similar results were obtained with Alr3423 (Fig. 3B, lanes 9 to 12), indicating that both kinases directly phosphorylate the All4160 STAS domain.

**Figure 3.**
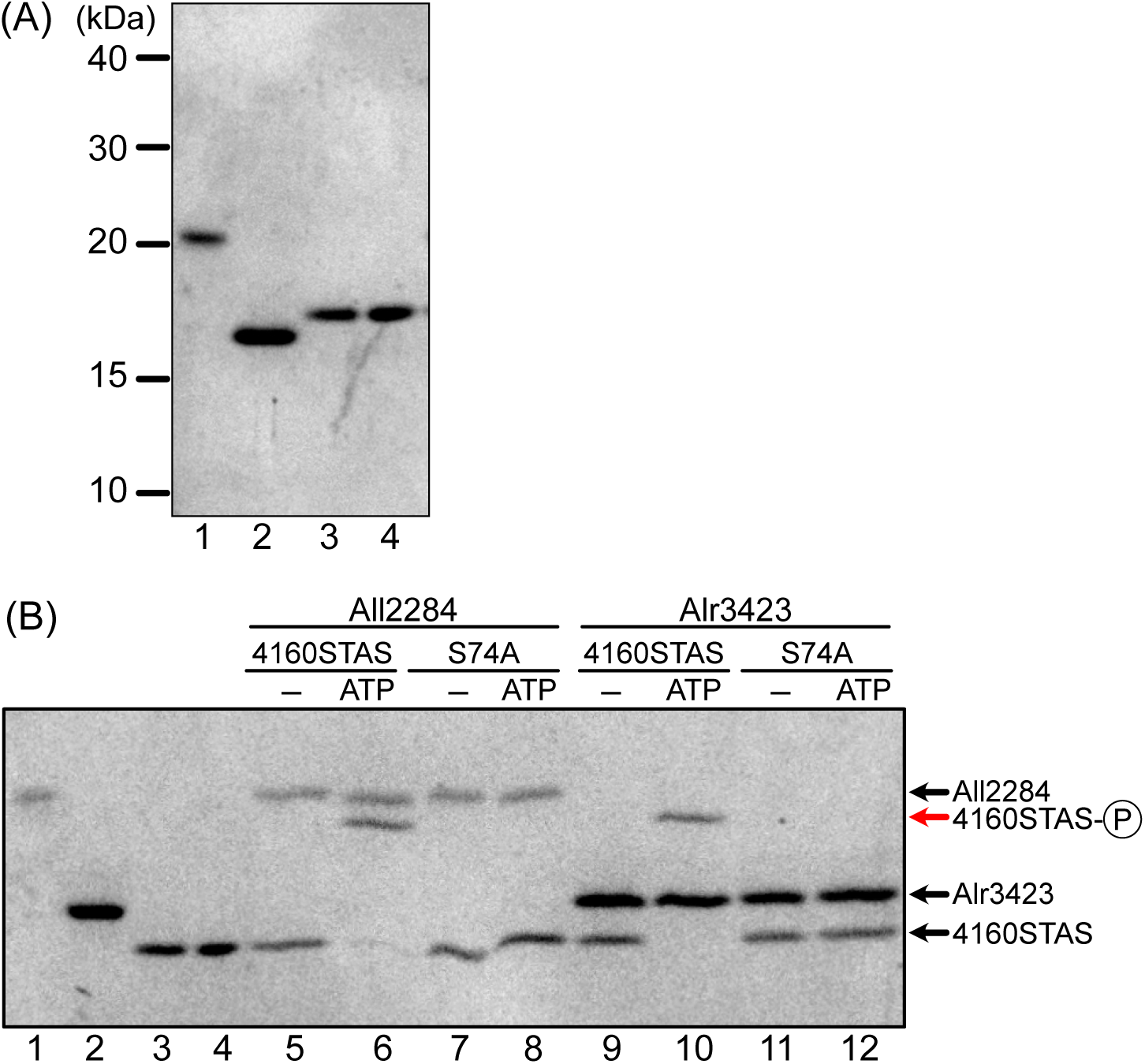
Phosphorylation of the All4160 STAS domain by All2284 and Alr3423 analyzed by Phos-tag SDS–PAGE. In vitro phosphorylation assays were performed using the All4160 STAS domain in the presence of All2284 or Alr3423. Both kinases phosphorylated the All4160 STAS domain in an ATP-dependent manner, as indicated by a mobility shift on Phos-tag SDS–PAGE. (A) Recombinant proteins used for the phosphorylation assay were analyzed by conventional SDS-PAGE. (B) In vitro phosphorylation assay analyzed by Phos-tag SDS-PAGE. Recombinant proteins were incubated with or without ATP prior to electrophoresis. Proteins included in each lane were as follows: lane 1, All2284; lane 2, Alr3423; lane 3, the All4160 STAS domain (4160STAS); lane 4, the S74A variant of the STAS domain (S74A); lanes 5–8, All2284 with 4160STAS or S74A; lanes 9–12, Alr3423 with 4160STAS or S74A. ATP was added to lanes 6, 8, 10, and 12.

### Genetic interactions of *henR* and candidate kinase genes

Both All2284 and Alr3423 were shown to phosphorylate the All4160 STAS domain in vitro (Fig. 3). We next investigated whether these kinases affect All4160 function in vivo. Because a nonphosphorylatable All4160 variant suppresses the defects of the *henR* mutant (Fig. 1), we hypothesized that loss of kinase activity responsible for All4160 phosphorylation would similarly suppress the *henR* mutant phenotype. To test this, we generated double mutants in which *henR* and either all2284 or alr3423 were disrupted. Single mutants of all2284 (Δ2284) and alr3423 (Δ3423) were able to grow under nitrogen-fixing conditions (Fig. 4). The Δ*henR*/Δ2284 double mutant exhibited impaired growth similar to that of Δ*henR*, whereas disruption of alr3423 in the Δ*henR* background restored diazotrophic growth (Fig. 4). These results indicate that Alr3423, but not All2284, is functionally involved in regulating All4160 activity during Hep layer formation.

**Figure 4.**
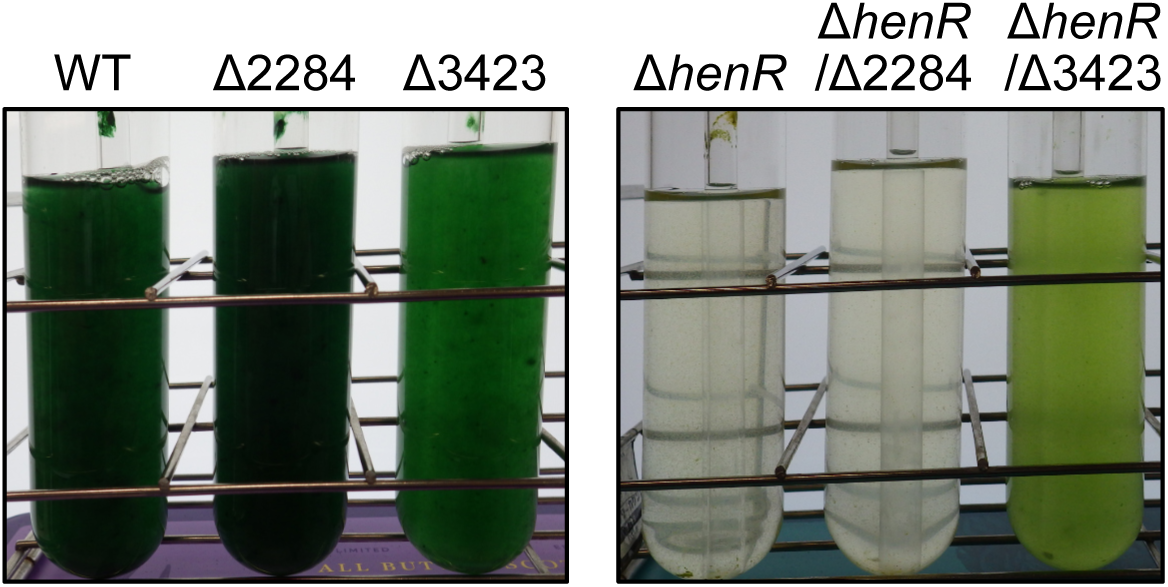
Genetic interaction between *henR* and candidate kinase genes all2284 and alr3423. Disruption of alr3423 restored diazotrophic growth of the *henR* mutant, whereas disruption of all2284 did not. Growth phenotypes of the indicated strains after 7 d of cultivation under nitrogen-fixing conditions. Representative culture tubes are shown. Strains included the wild type (WT), single mutants (Δ2284 and Δ3423), the *henR* mutant (Δ*henR*), and double mutants (Δ*henR*/Δ2284 and Δ*henR*/Δ3423).

## Discussion

In this study, we identified a phosphorylation-dependent regulatory module that controls heterocyst-specific polysaccharide (Hep) layer formation during heterocyst differentiation in *Anabaena*. The STAS domain-containing protein All4160 is negatively regulated by phosphorylation by Alr3423, and the SpoIIE-like phosphatase HenR is required for its activation (Fig. 5). Our results support a regulatory scheme in which All4160 activity is modulated through the antagonistic actions of a kinase and a phosphatase, reminiscent of a canonical partner-switching system. The observation that a nonphosphorylatable All4160 variant restores both Hep formation and diazotrophic growth in the absence of HenR (Fig. 1), together with the genetic interaction between *henR* and alr3423, in which disruption of alr3423 suppresses the defects of the *henR* mutant (Fig. 4), is consistent with this model. Although in vivo phosphorylation of All4160 remains to be directly demonstrated, the combined genetic and in vitro biochemical evidence strongly supports a phosphorylation-dependent regulatory mechanism controlling Hep layer formation.

**Figure 5.**
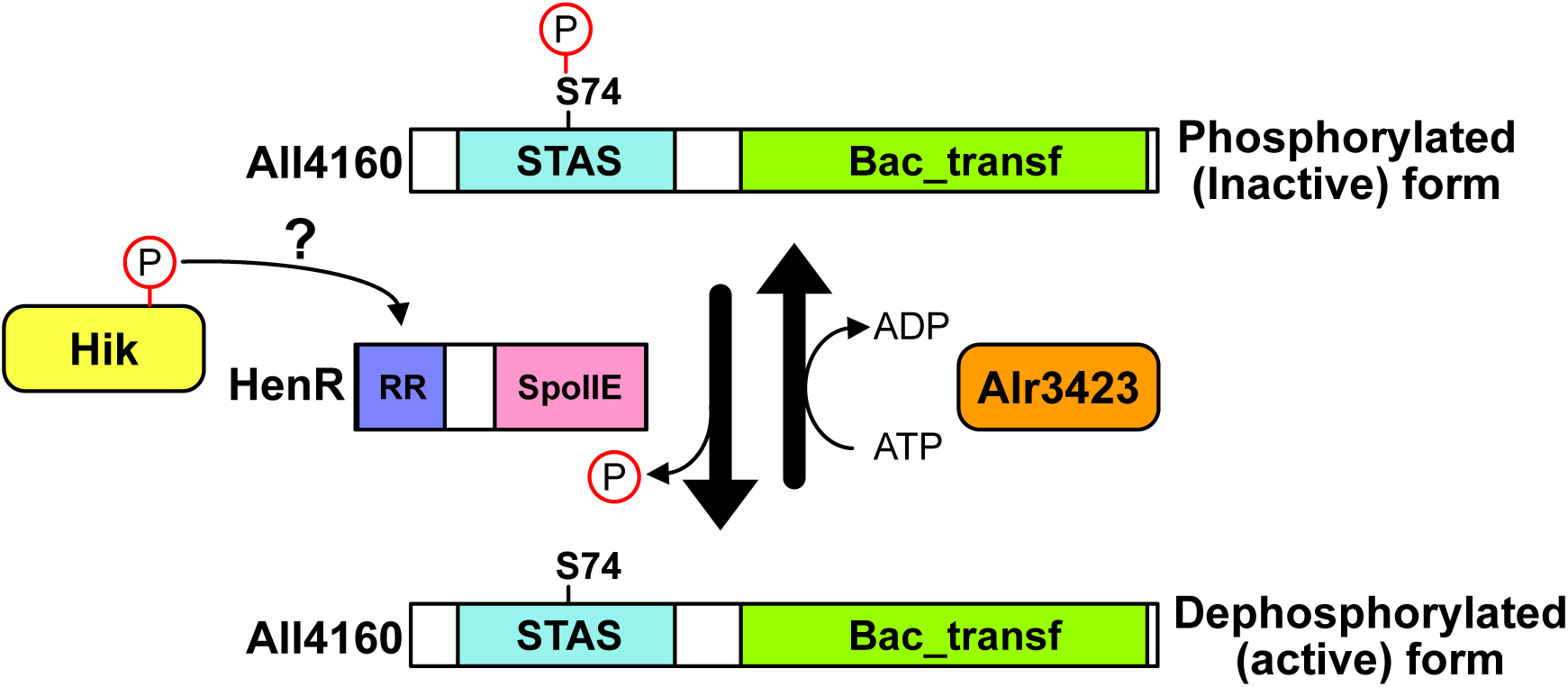
Phosphorylation-dependent regulation of All4160 activity. All4160 activity is suggested to be regulated by phosphorylation and dephosphorylation of its STAS domain, catalyzed by Alr3423 and HenR, respectively. HenR contains an N-terminal response regulator receiver (RR) domain and a C-terminal SpoIIE domain. HenR activity may be regulated by a cognate histidine kinase (Hik) as discussed in the text.

Notably, HenR contains an N-terminal receiver domain characteristic of response regulators, suggesting that its phosphatase activity may be subject to regulation by a cognate histidine kinase (Hik) (Fig. 5). This raises the possibility that Hep layer formation is controlled by modulation of HenR phosphatase activity through a hitherto unidentified Hik. Identification of the cognate Hik and the signals it perceives will be important for understanding the regulatory network governing Hep layer formation during heterocyst differentiation.

An additional unresolved question concerns the membrane topology and subcellular localization of All4160. Sequence analysis predicts a transmembrane domain between the STAS and glycosyltransferase domains, raising the possibility that these domains may reside on different sides of the membrane. Given that glycosyltransferase activity is generally expected to occur on the cytoplasmic side (15), this topology would place the STAS domain outside the cytoplasm, making it difficult to reconcile with regulation by cytoplasmic kinases and phosphatases. One possibility is that phosphorylation influences the membrane association or topology of All4160, thereby regulating access of the glycosyltransferase domain to its substrate. However, attempts to examine its localization using a GFP fusion were unsuccessful due to loss of protein function. Further studies will be required to clarify how phosphorylation affects the spatial organization and activity of All4160.

Previous studies have implicated PSS in EPS-related processes in diverse bacteria, including biofilm formation in *Vibrio fischeri* (16), symbiosis-associated polysaccharide production in *Sinorhizobium meliloti* (17), and EPS secretion required for hormogonium motility in *Nostoc punctiforme* (18). However, in these systems, the underlying mechanisms have not been fully elucidated, and it remains unclear whether PSS regulates EPS production through σ factor-dependent transcriptional control or through alternative mechanisms. Our findings provide a mechanistic framework for PSS involvement in polysaccharide biosynthesis by demonstrating direct regulation of a biosynthetic enzyme, rather than transcriptional control via σ factors. This mechanism may help explain how PSS contributes to diverse polysaccharide-related processes across phylogenetically distant bacteria.

All2284 was also identified as a kinase capable of phosphorylating the STAS domain of All4160 in vitro (Fig. 3); however, its disruption did not restore defects in diazotrophic growth in the Δ*henR* background (Fig. 4), indicating that All2284 is not involved in the phosphorylation-dependent regulatory module controlling Hep layer formation mediated by All4160 and HenR. The gene all2285, encoding a glycosyltransferase with an N-terminal STAS domain, is located adjacent to all2284 in the *Anabaena* genome (19). Given that All2285 is induced by desiccation and salt stress under the control of OrrA, and that exopolysaccharides (EPS) play important roles in desiccation tolerance in *Nostoc commune*, it is plausible that All2285 functions in EPS synthesis under these conditions (20, 21). In this context, All2284 may primarily act on All2285 rather than All4160. These observations raise the possibility that PSS-like regulatory modules have diversified to control distinct polysaccharide biosynthetic pathways in response to different environmental cues. Consistent with this idea, proteins containing both STAS and glycosyltransferase domains are widely distributed in cyanobacteria. For example, XssP, which is essential for the biosynthesis of the sulfated EPS synechan in *Synechocystis* sp. PCC 6803, is a homolog of All4160 (22). Synechan synthesis is known to respond to growth temperature, further suggesting that STAS-glycosyltransferase proteins may function in the regulation of environmentally responsive polysaccharide biosynthesis. The widespread occurrence of such STAS–glycosyltransferase proteins suggests that phosphorylation-dependent regulation of polysaccharide biosynthetic enzymes may represent a conserved regulatory strategy for controlling polysaccharide biosynthesis in cyanobacteria.

Several limitations of this study should be noted. First, phosphorylation of All4160 has not yet been directly demonstrated in vivo. Although we attempted to generate antibodies against the STAS domain of All4160, they did not allow reliable detection of the protein. Second, direct dephosphorylation of All4160 by HenR could not be demonstrated biochemically, because recombinant HenR could not be expressed in either *Escherichia coli* or insect cell expression systems. Third, the effect of phosphorylation on All4160 enzymatic activity has not yet been tested in vitro, because no activity assay is currently available owing to the unknown substrate of All4160.

Despite these limitations, multiple lines of genetic and biochemical evidence support the proposed model. In particular, the functional rescue by the nonphosphorylatable All4160 variant provides strong in vivo evidence that phosphorylation negatively regulates All4160 function. Future studies aimed at the direct detection of All4160 phosphorylation in vivo, biochemical reconstitution of HenR phosphatase activity, and the establishment of an assay system for All4160 activity will be important to further test and refine this model.

## Materials and Methods

### Strains and Growth conditions

*Anabaena* sp. strain PCC 7120 and its derivatives were grown in BG11 medium or BG11_0_ medium (BG11 lacking combined nitrogen) at 30°C under continuous illumination (30 µmol photons m^-2^ s^-1^) provided by cool white fluorescent light. Liquid cultures were bubbled with air supplemented with 1.0 % (v/v) CO_2_. For growth under nitrogen-fixing conditions, cells grown in BG11 were washed and resuspended in BG11_0_ as previously described (23). When required, spectinomycin and streptomycin were added at final concentrations of 2.5 µg ml[¹ each, and neomycin at 30 µg ml[¹.

### Mutant construction

All strains used in this study are listed in Table S1. Transformation of *Anabaena* was carried out by triparental conjugation as described by Elhai et al. (24). For construction of gene disruptants, the plasmid pRL271 (25), which carries *sacB*, was used, and antibiotic- and sucrose-resistant clones were selected to obtain double recombinants. Complete segregation was confirmed by PCR. To generate all4160-expressing strains in the Δ*henR* background, a DNA fragment containing the upstream region and coding sequence of all4160 was cloned into the suicide vector pSU101 (26), and single recombinants were selected based on neomycin resistance. A nonphosphorylatable variant of All4160 (S74A) was generated by site-directed mutagenesis, in which the nucleotides at positions 720 and 721 (A and G) were replaced with G and C, respectively, using the PrimeSTAR Mutagenesis Basal Kit (Takara Bio Inc.).

### Bacterial adenylate cyclase two-hybrid (BACTH) assay

Protein-protein interactions were analyzed using the bacterial adenylate cyclase two-hybrid system (Euromedex). A DNA fragment encoding the STAS domain (residues 1-139) of All4160 was amplified and cloned between the XbaI and SacI sites of pUT18C to generate an N-terminal fusion with the T18 fragment of adenylate cyclase. The S74A variant of the STAS domain was generated by site-directed mutagenesis. The coding regions of candidate kinases (All1702, All2284, Alr3423, Alr3655, and Alr3760) were amplified and cloned between the XbaI and EcoRI sites of pKT25 to generate N-terminal fusions with the T25 fragment. The resulting plasmids were co-transformed into *E*. *coli* BTH101. Transformants carrying both plasmids were grown in LB medium supplemented with 100 µg ml^-1^ ampicillin and 50 µg ml^-1^ kanamycin at 30°C. Aliquots (1 µl) of overnight cultures were spotted onto MacConkey agar plates supplemented with 1% (w/v) maltose, 0.5 mM IPTG, 100 µg ml^-1^ ampicillin and 50 µg ml^-1^ kanamycin, and incubated for 24 h at 30°C. Protein-protein interactions were evaluated based on colony color development compared with positive and negative controls.

### In vitro phosphorylation assay

His-tagged recombinant proteins were expressed from the expression vector pColdII (Takara Bio) in *E. coli* BL21(DE3) and purified using TALON Metal Affinity Resin (Takara Bio). To generate a plasmid for the STAS domain of All4160, a DNA fragment encoding the STAS domain (residues 1-148) of All4160 was amplified and cloned between the NdeI and EcoRI sites of pColdII. The S74A variant of the STAS domain was generated by site-directed mutagenesis. Plasmids for All2284 and Alr3423 were constructed by cloning the coding regions into the EcoRI and SalI sites of pColdII.

Purified STAS domain of All4160 (1 µg) or its S74A variant (1 µg) was incubated with All2284 (1 µg) or Alr3423 (1 µg) at 28°C for 30 min in phosphorylation buffer [25 mM Tris-HCl (pH 7.9), 50 mM NaCl, 5 mM MgCl_2_, and 1 mM dithiothreitol] in the presence or absence of 2 mM ATP. Reactions were terminated by the addition of 2x SDS-PAGE sample buffer, and proteins were resolved on 15% SDS-PAGE gels containing 75 µM Phos-tag acrylamide (FUJIFILM Wako Pure Chemical Corporation) and 150 µM MnCl_2_. After electrophoresis, proteins were visualized using EzStain Aqua (ATTO Corporation).

## Microscopy

Cells stained with Alcian blue were observed using a BX53 light microscope (Olympus). Bright-field images were acquired under transmitted light, and autofluorescence images were obtained using a U-FUW filter set.

## Acknowledgments

This work was supported by JSPS KAKENHI (Grant Numbers JP22K19138 and JP25K01938) and JST CREST (Grant Number JPMJCR25J3).

## Conflicts of Interest

The authors declare no conflicts of interest.

## Notes

### Competing Interest Statement

The authors have declared no competing interest.

## References

1. Berman-Frank I, Lundgren P, Falkowski P. 2003. Nitrogen fixation and photosynthetic oxygen evolution in cyanobacteria. Res Microbiol 154:157–164.

2. Flores E, Picossi S, Valladares A, Herrero A. 2019. Transcriptional regulation of development in heterocyst-forming cyanobacteria. Biochim Biophys Acta Gene Regul Mech 1862:673–684.

3. Kumar K, Mella-Herrera RA, Golden JW. 2010. Cyanobacterial heterocysts. Cold Spring Harb Perspect Biol2:a000315.

4. Huang G, Fan Q, Lechno-Yossef S, Wojciuch E, Wolk CP, Kaneko T, Tabata S. 2005. Clustered genes required for the synthesis of heterocyst envelope polysaccharide in *Anabaena* sp. strain PCC 7120. J Bacteriol 187:1114–1123.

5. Ehira S, Ohmori M, Sato N. 2003. Genome-wide expression analysis of the responses to nitrogen deprivation in the heterocyst-forming cyanobacterium *Anabaena* sp. strain PCC 7120. DNA Res 10:97–113.

6. Wang Y, Lechno-Yossef S, Gong Y, Fan Q, Wolk CP, Xu X. 2007. Predicted glycosyl transferase genes located outside the HEP island are required for formation of heterocyst envelope polysaccharide in *Anabaena* sp. strain PCC 7120. J Bacteriol 189:5372–5378.

7. Zhu J, Kong R, Wolk CP. 1998. Regulation of hepA of *Anabaena* sp. strain PCC 7120 by elements 5′ from the gene and by hepK. J Bacteriol 180:4233–4242.

8. Zhou R, Wolk CP. 2003. A two-component system mediates developmental regulation of biosynthesis of a heterocyst polysaccharide. J Biol Chem 278:19939–19946.

9. Ning D, Xu X. 2004. alr0117, a two-component histidine kinase gene, is involved in heterocyst development in *Anabaena* sp. PCC 7120. Microbiology 150:447–453.

10. Fan Q, Lechno-Yossef S, Ehira S, Kaneko T, Ohmori M, Sato N, Tabata S, Wolk CP. 2006. Signal transduction genes required for heterocyst maturation in *Anabaena* sp. strain PCC 7120. J Bacteriol 188:6688–6693.

11. Benson AK, Haldenwang WG. 1993. Bacillus subtilis σB is regulated by a binding protein (RsbW) that blocks its association with core RNA polymerase. Proc Natl Acad Sci U S A 90:2330–2334.

12. Duncan L, Losick R. 1993. SpoIIAB is an anti-sigma factor that binds to and inhibits transcription by regulatory protein sigma F from *Bacillus subtilis*. Proc Natl Acad Sci USA 90:2325–2329.

13. Bouillet S, Arabet D, Jourlin-Castelli C, Méjean V, Iobbi-Nivol C. 2018. Regulation of σ factors by conserved partner switches controlled by divergent signalling systems. Environ Microbiol Rep 10:127–139.

14. Yang X, Kang CM, Brody MS, Price CW. 1996. Opposing pairs of serine protein kinases and phosphatases transmit signals of environmental stress to activate a bacterial transcription factor. Genes Dev 10:2265–2275.

15. Schmid J, Sieber V, Rehm B. 2015. Bacterial exopolysaccharides: biosynthesis pathways and engineering strategies. Front Microbiol 6:1–24.

16. Morris AR, Visick KL. 2013. The response regulator SypE controls biofilm formation and colonization through phosphorylation of the syp-encoded regulator SypA in *Vibrio fischeri*. Mol Microbiol 87:509–525.

17. Baena I, Pérez-Mendoza D, Sauviac L, Francesch K, Martín M, Rivilla R, Bonilla I, Bruand C, Sanjuán J, Lloret J. 2019. A partner-switching system controls activation of mixed-linkage β-glucan synthesis by c-di-GMP in *Sinorhizobium meliloti*. Environ Microbiol 21:3379–3391.

18. Riley KW, Gonzalez A, Risser DD. 2018. A partner-switching regulatory system controls hormogonium development in the filamentous cyanobacterium *Nostoc punctiforme*. Mol Microbiol 109:555–569.

19. Kaneko T, Nakamura Y, Wolk CP, et al. 2001. Complete genomic sequence of the filamentous nitrogen-fixing cyanobacterium *Anabaena* sp. strain PCC 7120. DNA Res 8:205–213, 227–253.

20. Tamaru Y, Takani Y, Yoshida T, Sakamoto T. 2005. Crucial role of extracellular polysaccharides in desiccation and freezing tolerance in the terrestrial cyanobacterium *Nostoc commune*. Appl Environ Microbiol 71:7327–7333.

21. Kimura S, Sato M, Fan X, Ohmori M, Ehira S. 2022. The two-component response regulator OrrA confers dehydration tolerance by regulating avaKa expression in the cyanobacterium *Anabaena* sp. strain PCC 7120. Environ Microbiol 24:5165–5173.

22. Maeda K, Okuda Y, Enomoto G, Watanabe S, Ikeuchi M. 2021. Biosynthesis of a sulfated exopolysaccharide, synechan, and bloom formation in the model cyanobacterium *Synechocystis* sp. strain PCC 6803. eLife10:e66538.

23. Ehira S, Ohmori M. 2006. NrrA, a nitrogen-responsive response regulator facilitates heterocyst development in the cyanobacterium *Anabaena* sp. strain PCC 7120. Mol Microbiol 59:1692–1703.

24. Elhai J, Vepritskiy A, Muro-Pastor AM, Flores E, Wolk CP. 1997. Reduction of conjugal transfer efficiency by three restriction activities of *Anabaena* sp. strain PCC 7120. J Bacteriol 179:1998–2005.

25. Black TA, Cai Y, Wolk CP. 1993. Spatial expression and autoregulation of hetR, a gene involved in the control of heterocyst development in *Anabaena*. Mol Microbiol 9:77–84.

26. Ehira S, Shimmori Y, Watanabe S, Kato H, Yoshikawa H, Ohmori M. 2017. The nitrogen-regulated response regulator NrrA is a conserved regulator of glycogen catabolism in β-cyanobacteria. Microbiology 163:1711–1719.

