## Supplemental Data for "A phosphorylation-dependent partner-switching-like module controls heterocyst polysaccharide layer formation in *Anabaena* sp. strain PCC 7120"

Table S1. Strains used in this study

| Strains | Description |
|---|---|
| WT | *Anabaena* sp. strain PCC 7120 |
| Δ *henR* | The *henR* coding region (1,137 bp) was disrupted by replacing nucleotides 128–1,129 with a spectinomycin/streptomycin resistance cassette derived from pDW9 (1) in WT. |
| Δ 4160 | The *all4160* coding region (1,050 bp) was disrupted by replacing nucleotides 53–897 with a neomycin resistance cassette derived from pRL161 (2) in WT. |
| Δ *henR*/all4160 | The plasmid pSU101 (3), carrying the upstream region (1,170 bp), coding sequence, and downstream region (42 bp) of *all4160*, was integrated into the all4160 locus of the Δ *henR* strain by homologous recombination via a single-crossover event. |
| Δ *henR*/S74A | Same as the Δ *henR*/all4160 strain, except that Ser74 of All4160 was substituted with Ala. |
| Δ 2284 | The *all2284* coding region (498 bp) was disrupted by replacing nucleotides 25–384 with a neomycin resistance cassette derived from pRL161 (2) in WT. |
| Δ 3423 | The *alr3423* coding region (444 bp) was disrupted by replacing nucleotides 6–380 with a neomycin resistance cassette derived from pRL161 (2) in WT. |
| Δ *henR*/ Δ 2284 | The *all2284* gene was disrupted in the Δ *henR* strain. |
| Δ *henR*/ Δ 3423 | The *alr3423* gene was disrupted in the Δ *henR* strain. |

References


1. **Golden JW**, **Wiest DR**. 1988. Genome Rearrangement and Nitrogen Fixation in Anabaena Blocked by Inactivation of xisA Gene. Science **242**:1421–1423.

2. **Elhai J**, **Wolk CP**. 1988. A versatile class of positive-selection vectors based on the nonviability of palindrome-containing plasmids that allows cloning into long polylinkers. Gene **68**:119–138.

3. **Ehira S**, **Shimmori Y**, **Watanabe S**, **Kato H**, **Yoshikawa H**, **Ohmori M**. 2017. The nitrogen-regulated response regulator NrrA is a conserved regulator of glycogen catabolism in β-cyanobacteria. Microbiology **163**:1711–1719.

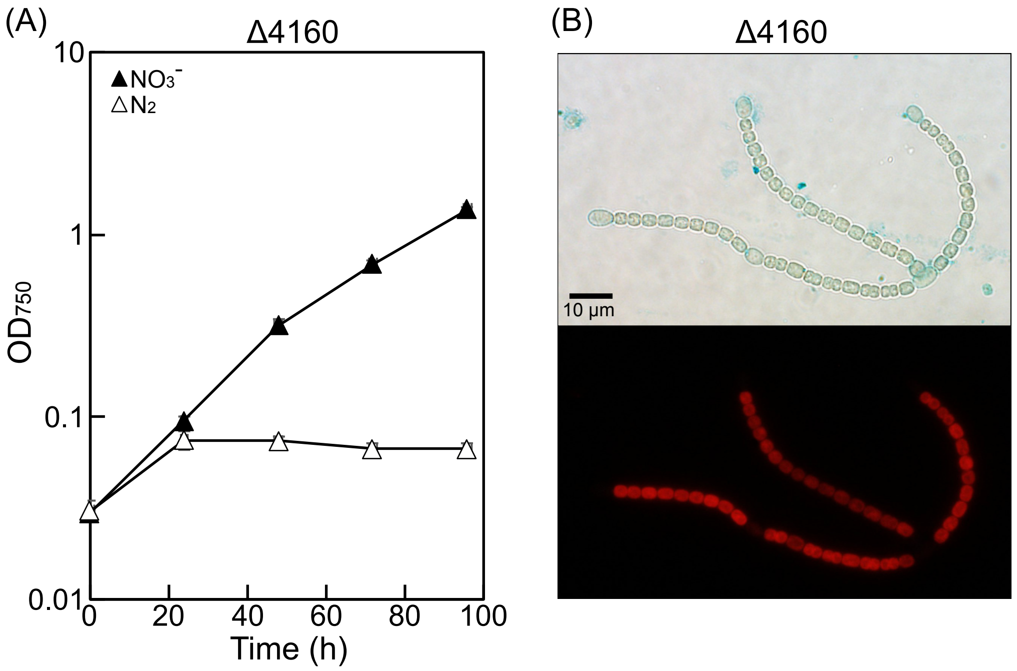


FigureS1. Phenotypes of the all4160 mutant (Δ4160)

(A) Growth of Δ4160 with nitrate (closed triangles) or dinitrogen (open triangles) as nitrogen sources.

(B) Micrographs of Alcian blue-stained Δ4160 cells. The upper panel shows bright-field images, and the lower panel shows autofluorescence.

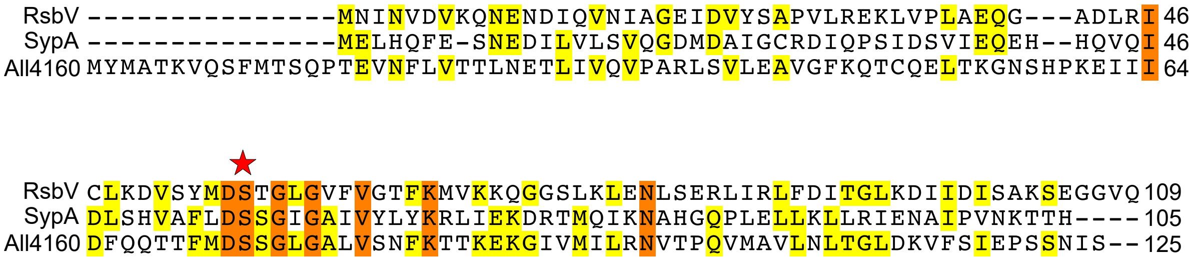


Figure S2. Multiple sequence alignment of STAS domain amino acid sequences.

A multiple alignment was generated using the STAS domains of RsbV from *Bacillus subtilis*, SypA from *Vibrio fischeri*, and All4160. Serine residues known to be phosphorylated in RsbV and SypA are indicated by an asterisk above the alignment (Yang *et al*., 1996; Morris and Visick, 2013). The corresponding residue in All4160 (Ser74) is conserved.

### References


1. Morris AR, Visick KL. 2013. The response regulator SypE controls biofilm formation and colonization through phosphorylation of the syp-encoded regulator SypA in Vibrio fischeri. Mol. Microbiol. 87:509–25.

2. Yang X, Kang CM, Brody MS, Price CW. 1996. Opposing pairs of serine protein kinases and phosphatases transmit signals of environmental stress to activate a bacterial transcription factor. Genes Dev. 10:2265–2275.

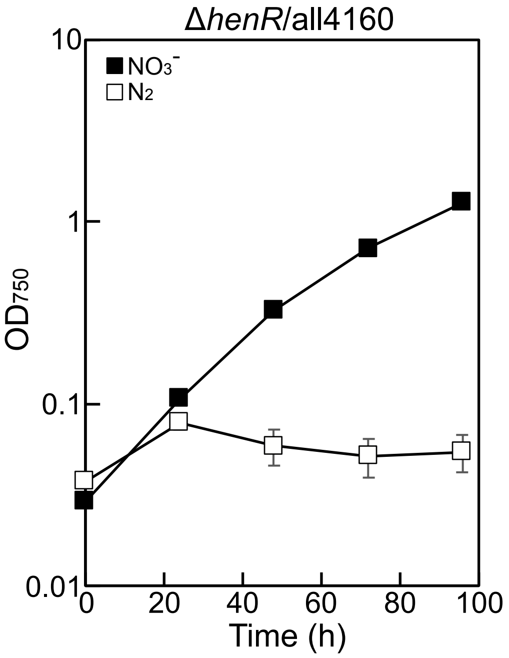


Figure S3. Expression of wild-type All4160 does not restore the nitrogen-fixation defect of Δ*henR*.

Growth of Δ*henR* carrying a plasmid expressing wild-type All4160 with nitrate (closed squares) or dinitrogen (open squares) as nitrogen sources.

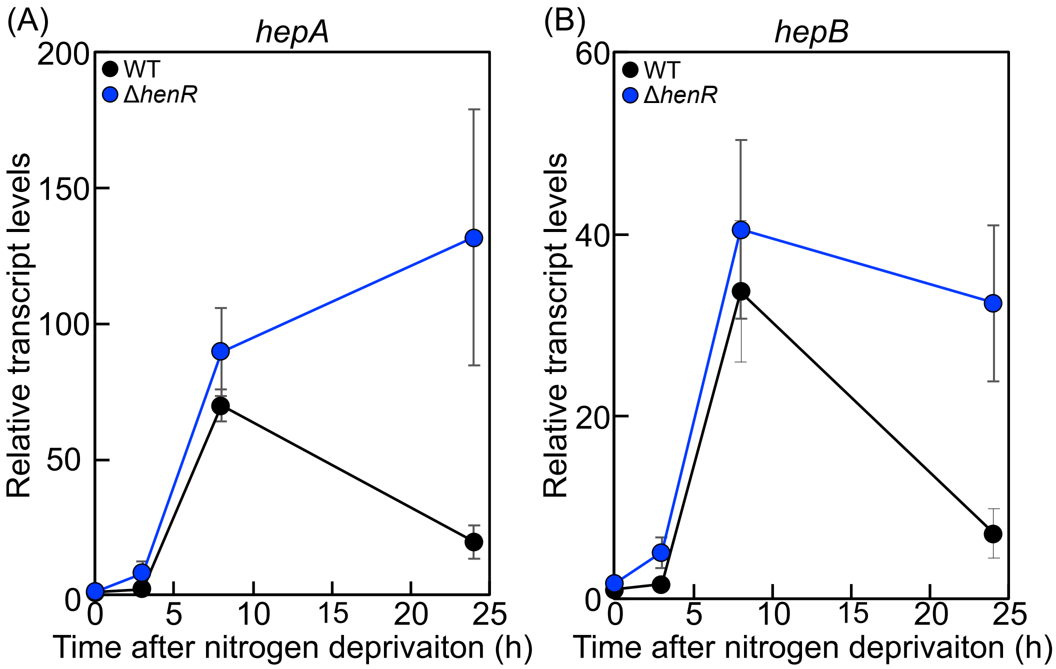


Figure S4. Disruption of *henR* does not affect transcription of *hep* genes.

Transcript levels of *hepA* (A) and *hepB* (B) after nitrogen deprivation were determined by qRT-PCR in the wild-type strain (WT) (closed circles) and Δ*henR* (blue circles). Transcript levels were normalized to the WT value at 0 h, which was set to 1.
